# Edge-centric functional network representations of human cerebral cortex reveal overlapping system-level architecture

**DOI:** 10.1101/799924

**Authors:** Joshua Faskowitz, Farnaz Zamani Esfahlani, Youngheun Jo, Olaf Sporns, Richard F. Betzel

## Abstract

Network neuroscience has relied on a node-centric network model in which cells, populations, and regions are linked to one another *via* anatomical or functional connections. This model cannot account for interactions of edges with one another. Here, we develop an *edge*-centric network model, which generates the novel constructs of “edge time series” and “edge functional connectivity” (eFC). Using network analysis, we show that at rest eFC is consistent across datasets and reproducible within the same individual over multiple scan sessions. We demonstrate that clustering eFC yields communities of edges that naturally divide the brain into overlapping clusters, with regions in sensorimotor and attentional networks exhibiting the greatest levels of overlap. We go on to show that eFC is systematically and consistently modulated by variation in sensory input. In future work, the edge-centric approach could be used to map the connectional architecture of brain circuits and for the development of brain-based biomarkers of disease and development.

## INTRODUCTION

Network science offers a promising framework for representing and modeling neural systems across many spatial scales [1]. From interconnected cells [2], to neuronal populations [3, 4], to large-scale brain areas [5, 6], network analysis has contributed insight into the topological principles that govern nervous system organization and shape brain function. These include small-world architecture [7], the emergence of integrative hubs and rich clubs [8–10], modular structure to promote specialized information processing [11–13], and tradeoffs between these features with the material and metabolic costs of wiring the brain together [14, 15].

Central to these and other discoveries in network neuroscience is a simple representation of the brain in which neural elements and their pairwise interactions are treated as the nodes and edges of a network, respectively [16, 17]. This standard model is fundamentally node-centric in that it treats neural elements (the nodes) as the irreducible units of brain structure and function. This emphasis on network nodes is further reinforced by the types of analyses carried out on brain networks, which tend to focus on properties of nodes or groups of nodes, e.g. their centralities, community affiliations, etc. [18].

A limitation of the node-centric approach is that it cannot capture potentially meaningful features or patterns of interrelationships among edges. In other scientific domains, prioritizing network edges, for example by modeling and analyzing edge-edge interactions as a graph, has provided important insights into the organization and function of complex systems [19–22]. In contrast, with few exceptions [23], network neuroscience has remained focused on nodal features and partitions, paralleling a rich history of parceling, mapping, and comparing cortical and subcortical gray matter regions [24].

Here, we present a novel modeling framework for investigating functional brain network data from an *edge*-centric perspective. Our approach can be viewed as a temporal “unwrapping” of the Pearson correlation measure – the metric commonly used for estimating the strength of functional connectivity between pairs of brain regions [25] – thereby generating interpretable time series for each edge that express fluctuations in its weight across time.

Importantly, edge time series allow the estimation of edge correlation structure, a construct we refer to as edge functional connectivity (eFC). High eFC indexes strong similarity in the co-fluctuation of two edges across time, while low eFC indicates co-fluctuation patterns that are largely independent. Here, we first demonstrate that eFC is highly replicable given 30 minutes worth of data, is stable within individuals across multiple scan sessions, and is consistent across datasets. Next, we applied data-driven clustering algorithms to eFC, which resulted in partitions of the eFC network into communities of co-fluctuating edges. Each community could be mapped back to individual nodes, yielding overlapping regional community assignments. We found that some nodes were affiliated with relatively few communities, while nodes associated with sensorimotor and attentional networks participated in disproportionately many communities. Finally, we compared the organization of eFC at rest and during passive viewing of movies, and found that eFC was consistently and reliably modulated by changes in sensory input.

## RESULTS

In this section we analyze edge functional connectivity (eFC) estimated using neuroimaging data from three independently acquired datasets: 100 unrelated subjects from the Human Connectome Project (HCP; [26]), ten subjects scanned ten times as part of the Midnight Scan Club (MSC; [27]), and ten subjects scanned multiple times as part of the Healthy Brain Network Serial Scanning Initiative (HBN; [28]).

### Edge functional connectivity

Many studies have investigated networks whose nodes and edges represent brain regions and pairwise functional interactions, respectively [5, 29]. Here, we extend this framework to consider interactions not between pairs of brain regions, but pairs of *edges*.

Extant approaches for estimating edge-edge connectivity matrices include construction of line graphs [20] or calculating edge overlap indices [19]. While suitable for sparse networks with positively-weighted edges, these approaches are less appropriate for functional neuroimaging data, where networks are typically fully-weighted and signed. Here, we introduce a method that is well-suited for these types of data, operates directly on time series, and is closely related to the Pearson correlation coefficient (typically used to assess strength of inter-regional functional connections). We refer to the matrices obtained using this procedure as “edge functional connectivity” (eFC).

Beginning with regional time series, calculating eFC can be accomplished in three steps, starting by z-scoring the time series (Fig. 1a,d). Next, for all pairs of brain regions, we calculate the element-wise product of their z-scored time series (Fig. 1b,d). This results in a new set of time series, referred to as “edge time series,” whose elements represent the instantaneous co-fluctuation magnitude for pairs of brain regions and whose average across time is equal to the Pearson correlation coefficient (Fig. 1c). These values are positive when activity of two regions fluctuates in the same direction at precisely the same moment in time (although we note that this co-fluctuation time series could easily be calculated at non-zero lags, as well), are negative when activity fluctuates in opposite directions, and zero when activity fluctuates close to baseline. Importantly, the magnitude of these edge time series are not systematically related to in-scanner motion (Fig. S1). The third and final step involves calculating the scalar product between pairs of edge time series. When repeated over all pairs of edges, the result is an edge-by-edge matrix, whose elements we normalize to the interval [−1, 1] (Fig. 1f and Fig. 2a). See **Materials and Methods** for additional details on eFC construction.

**FIG. 1.**
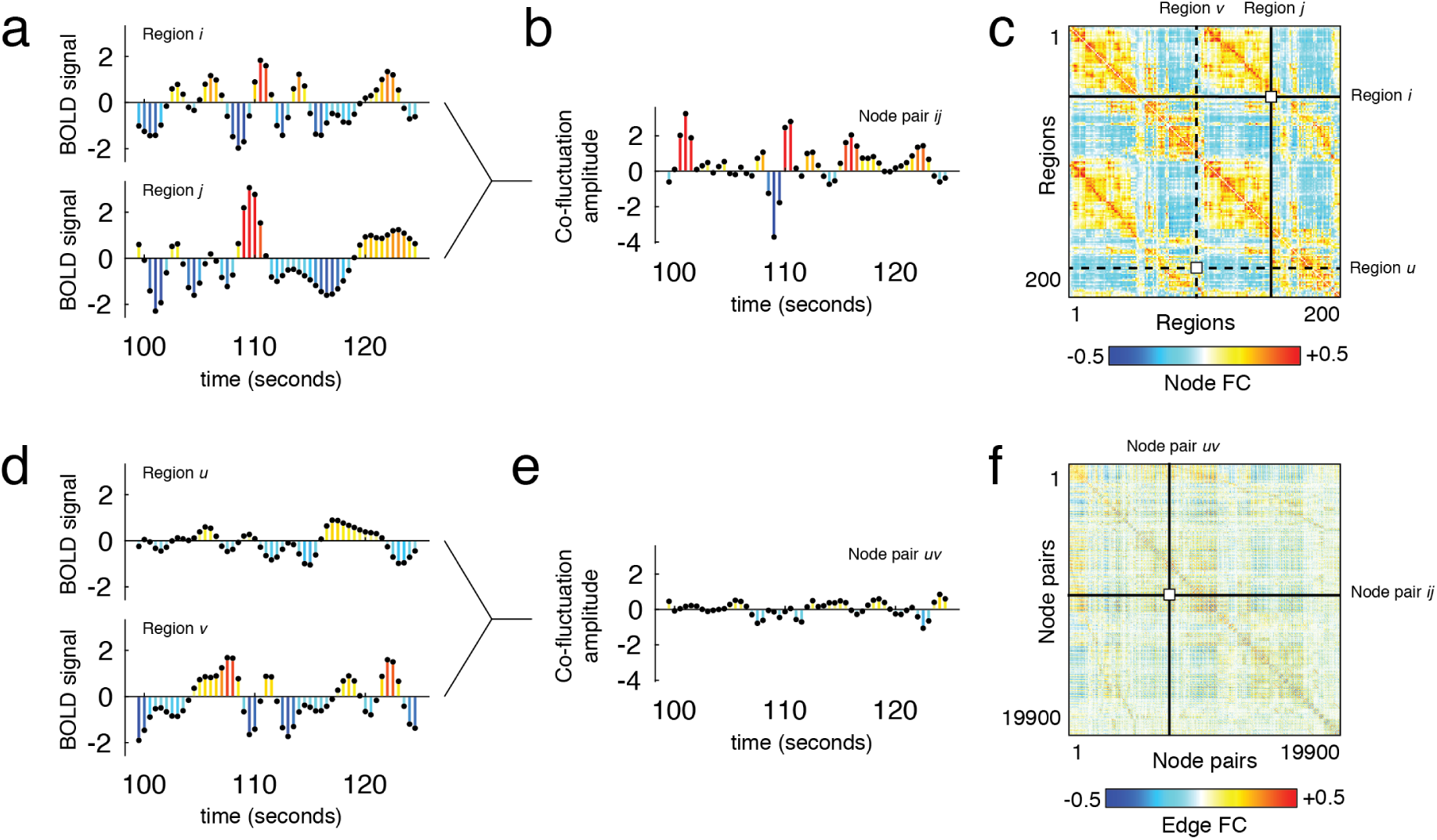
Derivation of edge functional connectivity (eFC) matrix. Each element of the eFC matrix is calculated based on the fMRI BOLD activity time series from four nodes (brain regions). In panels *a* and *d*, we show four representative times series from regions *i, h, u*, and *v*. Nodal FC (panel *c*; nFC) is typically calculated by standardizing (z-scoring) each time series, performing an element-wise product (dot product) of time series pairs, and calculating the average value of a product time series (actually the sum of each element divided by *T* − 1, where *T* is the number of elements or observations in the time series). To calculate eFC, we retain the vectors of element-wise products for every pair of regions. In panels *b* and *e* we show product time series for the pairs {*i, j*} and {*u, v*}, respectively. The elements of these product time series represent the magnitude of time-resolved co-fluctuation between region pairs (or edges in the nFC graph). We can calculate the magnitude of eFC by performing an element-wise multiplication of the product time series and normalizing the sum by the squared root standard deviations of both time series, ensuring that the magnitude of eFC is bounded to the interval [-1, 1]. The resulting value is stored in the eFC matrix, shown in panel *f*.

**FIG. 2.**
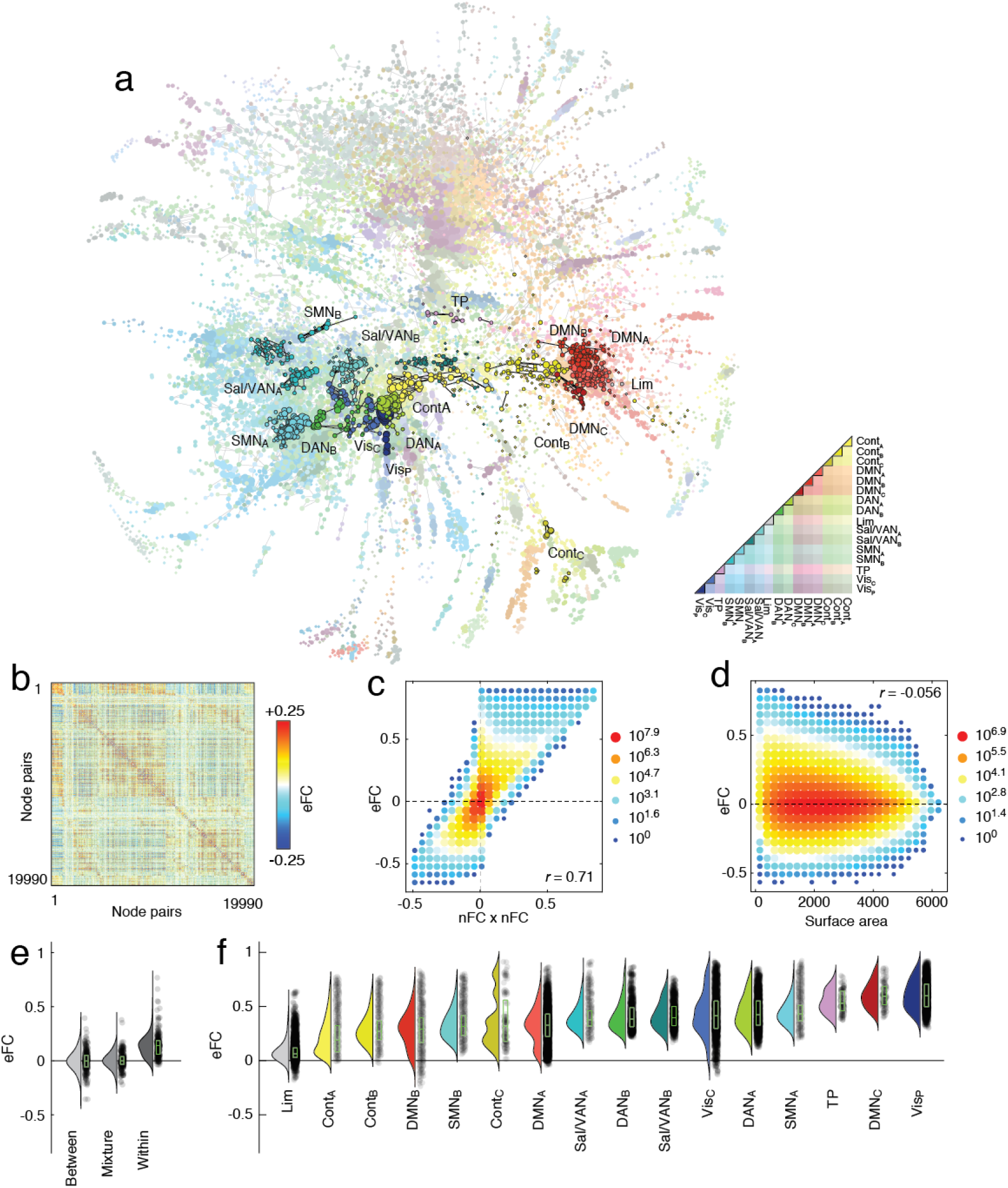
Organization of edge functional connectivity (eFC). (*a*) Force-directed layout of eFC matrix (largest connected component). Nodes in this graph represent edges in the traditional nodal functional connectivity (nFC) matrix. Here, nodes are colored according to whether the corresponding edge fell within or between cognitive systems. Within-system edges are encircled in black. (*b*) eFC matrix in which rows and columns correspond to pairs of brain regions (see Fig. 1 for details of how this matrix was generated). (*c*) Two-dimensional histogram of relationship between eFC and the product of edges’ respective nFC weights. (*d*) Two-dimensional histogram of relationship between eFC and the surface area of the quadrilateral defined by the four nodes. (*e*) Mean eFC among edges where both edges fall between systems (between), where one edge falls within and the other between systems (mixture), and where both edges fall within systems (within). (*f*) Mean eFC among edges within sixteen cognitive systems.

While eFC is, to our knowledge, a novel construct, we note that the first two steps in calculating eFC are the same as those used to calculate nodal FC (nFC); the mean value of any co-fluctuation time series is simply the Pearson correlation coefficient. Whereas subsequent sections investigate the organization of eFC, the remainder of this section is spent grounding eFC by comparing it with more familiar measurements.

Given that eFC is mathematically related to nFC, we first asked whether it was possible to approximate eFC using estimates of nFC. This is an important question; while the calculation of eFC can be implemented efficiently, performing certain operations on the eFC matrix can prove computationally expensive (it is a fully-weighted [*N*(*N* − 1)/2 × *N* (*N* − 1)/2] matrix, where *N* is the number of brain regions; Fig. 2b). Perhaps the simplest means of comparison is to relate the weight of the edge connection between region pairs {*i, j*} and {*u, v*} with the product of those edges in the nFC matrix. Essentially this comparison assesses whether eFC contains additional information above and beyond nFC. We found that, as expected, these two measures were correlated (*r* = 0.71; *p <* 10^−15^; Fig. 2c), but were nonetheless not identical. This observation indicates that, while there exists some shared variance, eFC contains information not obviously accounted for by nFC. We note, also, that the weights of eFC are similar across three independently acquired datasets (Fig. S2) and irrespective of whether or not we performed global signal regression (Fig. S3).

Next, we asked whether eFC exhibits any clear spatial dependence, as nFC is known to decay as a function of Euclidean distance [30, 31]. This requires computing a spatial relationship between two pairs of brain regions; hence the traditional metric of Euclidean distance is insufficient. Instead, we assessed the spatial dispersion of eFC with the surface area of the quadrilateral formed by the centroids of the brain region pairs. We found evidence of a weak negative relationship between surface area and eFC (*r* = −0.056, *p <* 10^−15^; Fig. 2d), suggesting that unlike traditional nFC, whose connection weights are more strongly influenced by spatial relationships of brain areas to one another [30–32], eFC is less constrained by the brain’s geometry.

Finally, we asked whether eFC bears the imprint of nFC communities – brain regions whose activity is highly correlated with members of its own community, but weakly or anti-correlated with members of other communities [6, 11–13]. We asked whether the magnitude of eFC of within-community connections was distinguishable from that of between-community connections. To address this question we classified every edge in the nFC network according to whether it fell within or between communities [33], resulting in three possible combinations of connections in the eFC graph: eFC could link two edges that both fell within a community, two edges that both fell between communities, or an edge that fell within to an edge that fell between. In general, we found that eFC was significantly stronger for within-community edges compared to the other two categories (Fig. 2e). Interestingly, we found eFC could be distinguished further by dividing within-community edges by cognitive system [33] (Fig. 2f).

Collectively, these results suggest that, while eFC is related to existing concepts previously applied to nFC, eFC also caputures unique signal components and exhibits unique architectural features above and beyond those observed in nFC. These observations motivate further exploration of eFC and its organization as a network.

### Edge functional connectivity **is** stable within individuals

In the previous section we described the procedure for calculating eFC and related it to several well-known properties of nFC. In this section, we test the robustness of eFC to scan duration, i.e. how much data is required before eFC stabilizes, and also test whether eFC is stable across repeated scans of the same individual.

To address these questions we leveraged the within-subject design of the MSC dataset. For each subject, we aggregated their fMRI data across all scan sessions and estimated a single eFC matrix. Then, we randomly and uniformly sampled smaller amounts of data (25 samples per subject) and estimated eFC, which we compared against the aggregated eFC matrix. Similar to other studies [34], we found that with small amounts of data eFC was highly variable (Fig. 3a). However, we observed a monotonic increase in similarity as a function of time, so that with 30 minutes of data, the similarity was much greater (*r* = 0.88). This is of practical significance; like traditional nFC, it implies that eFC estimated using data from short scan sessions may not deliver accurate representations of an individuals’ edge network organization.

**FIG. 3.**
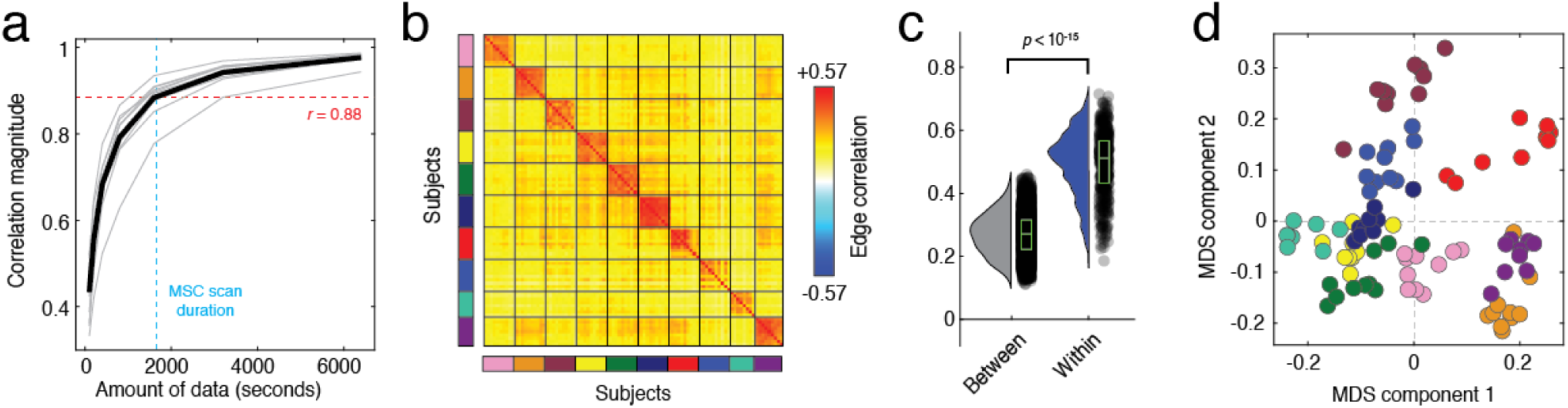
Intra- and inter-subject similarity of eFC across scan sessions. All panels from this figure were generated using data from the Midnight Scan Club. (*a*) Correlation session-averaged eFC matrices with eFC estimated using different amounts of data; mean value shown as black line. (*b*) Similarity of eFC within and between subjects. Each block corresponds to data from a single subject; subjects are also identifiable by the color of the rectangle alongside each bock. (*c*) Violin plots of within- and between-subject similarity values from the matrix shown in panel *b*. (*d*) Scan sessions plotted according to coordinates generated by performing a two-dimensional multi-dimensional scaling analysis of the matrix in panel *b*. Note that scans from the same subject (shown here with the same color) are located near each other.

Next, we examined the reliability of eFC over multiple scan sessions. That is, if we imaged an individual on separate days, would their eFC on those days be more similar to each other than to that of a different individual? The MSC dataset is well-poised to address this question; in the MSC dataset ten individuals were imaged ten times (30 minute resting-state runs). For all subjects and for all datasets, we estimated eFC and calculated the pairwise similarity (Pearson correlation) between all pairs of subjects and scans. We found eFC to be highly reliable in the MSC dataset, where the mean within-subject similarity was *r* = 0.50 ± 0.10 compared to *r* = 0.27 ± 0.07 between subjects (Fig. 3b,c). In Fig. 3d, we show the results of applying multi-dimensional scaling to the similarity matrix from Fig. 3b. Note that scans from the same subject, distinguished by color, are located near one another in this low-dimensional approximation. We found similar results in the HBN and HCP datasets (see Fig. S4).

Collectively, these findings suggest that eFC is a reliable marker of an individual, provided that it was estimated using enough data. This observation serves as an important validation of eFC, and suggests that eFC may be useful in future applications as substrate for biomarker generation [35] and “fingerprinting” [36], i.e. mapping the connectional features that can accurately distinguish individuals from one another versus those features that are shared across individuals.

### The overlapping community structure of human cerebral cortex

The previous two sections related eFC to existing concepts based on nFC and showed that it can be a reliable marker of an individual’s brain organization. In this section, we show that eFC can provide novel neuroscientific insight. Specifically, we leverage eFC to partition brain regions into overlapping communities.

While many studies have investigated the brain’s community structure [6, 11, 13, 37–39], most have relied on methodology that forces each brain region into one and only one community. However, partitioning brain regions into non-overlapping communities clashes with evidence suggesting that any given aspect of cognition and behavior requires contributions from regions that span multiple nodally-defined communities and systems [40, 41]. Non-overlapping nodal partitions are hence inconsistent with the notion that functional contributions of brain regions are massively overlapping, with regions becoming dynamically engaged and recruited into functional circuits with shifting demands imposed by varying internal and external states [42]. Accordingly, a more flexible definition of communities that matches more closely the multifunctional nature of brain regions is needed [43].

One strategy is to derive overlapping communities, allowing brain regions to be affiliated with multiple communities at the same time. While deriving overlapping communities can be challenging when using nFC, doing so using eFC is straightforward. Clustering the eFC graph assigns each edge in the network to a nonoverlapping edge community. Each edge is also associated with two brain regions (the nodes it connects). Thus edge community assignments can be mapped back onto individual brain regions and because every region is associated with *N* − 1 edges, each of which can maintain a different community assignment, every region can have multiple community affiliations.

In this section, we cluster eFC matrices to discover overlapping communities in human cerebral cortex. More specifically, we use a modified *k*-means to partition the eFC graph into non-overlapping communities and map the edge assignments back to individual nodes. We note that, in general, other community detection algorithms could be used in place of *k*-means; our decision to use this algorithm was practically motivated, as *k*-means exhibited significantly faster runtimes than other algorithms, e.g. modularity maximization [44] and Infomap [45], which have been used extensively in previous work to derive communities in both functional and structural brain networks.

In Fig. 4 we show representative communities obtained with *k* = 10 (See Fig. S6 for examples with different numbers of communities). To demonstrate that the communities capture meaningful variance in our data, we show the edge co-fluctuation time series, the eFC matrix, and the community co-assignment matrix reordered according to the derived communities (Fig. 4a,b,c). Here, the elements of the co-assignment matrix represent the probability that two edges were assigned to the same community across partitions as we varied the number of communities from *k* = 2 to *k* = 20.

**FIG. 4.**
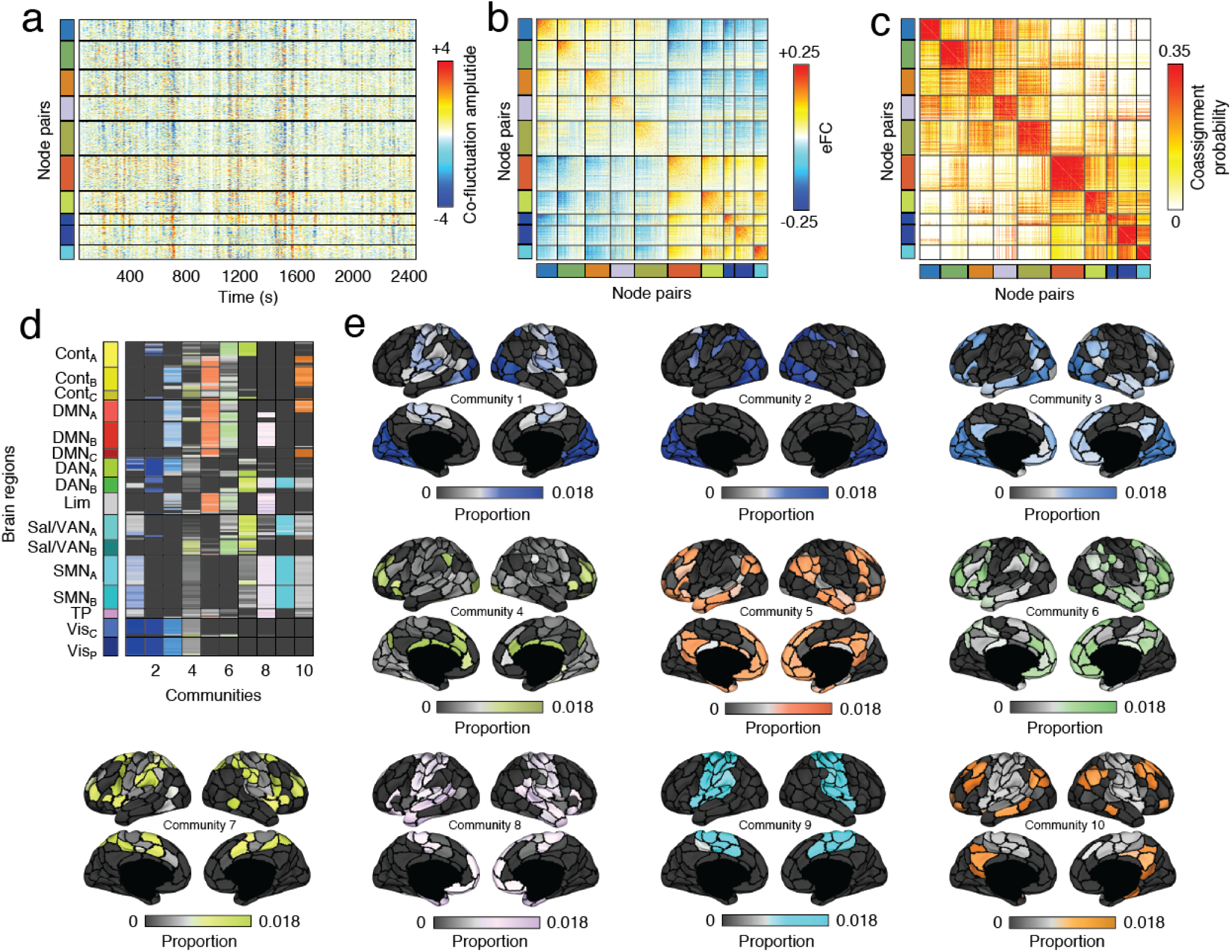
Edge communities. We applied similarity-based clustering to eFC from the HCP dataset. Here, we show results with the number of clusters fixed at *k* = 10. (*a*) Here, we reordered edge time series according to the detected community assignments. In this panel, horizontal lines divide communities from each other. The colors to the left of the time series plots identify each of the ten communities. (*b*) We also reordered the rows and columns of the eFC matrix to highlight the same ten communities. Note that, on average, within-community eFC is greater than between-community eFC. (*c*) We calculated the probability that pairs of edges (node pairs) were co-assigned to the same community. Here, we show the co-assignment matrix with rows and columned reordered according to community assignments. Note that, in general, the range of co-assignment probabilities goes to 1. Here, we truncate the color limits to emphasize the 10-community partition (In Fig. S5 we show the same co-assignment matrix at different values of *k* and with non-truncated color limits). We present two visualizations of the edge communities projected back to brain regions (nodes). In *d*, we depict overlapping communities in matrix form. Each column in this matrix represents one of ten communities. For each community and for each node, we calculated the proportion of all edges assigned to that community that included that node as one of its endpoints (“stubs”), indicated by the color and brightness of each cell. Dark colors indicate few edges; bright colors indicate many (*e*) Topographic distribution of communities. Note that many of the communities resemble known intrinsic connectivity networks. However, because the communities here are allowed to overlap, it is possible for nodes associated with a particular intrinsic connectivity network to participate in multiple edge communities.

While the communities detected here are defined at the level of edges rather than nodes, they naturally projectedge communities back onto brain regions. This was accomplished by extracting the edges associated with each community, determining which nodes were at the endpoints of each edge (the “stubs”), and counting the number of times that each node was represented in this stub list. We show these results in matrix form in Fig. 4d. In this panel, rows and columns represent nodes and edge communities, with nodes ordered according to the canonical system labels described in [33]. Bright-colored cells indicate that the corresponding node contributes many of its edges to a given community, while dark-colored cells indicate that the node contributes few or none of its edges. Note that, because a node’s edges can be assigned to different communities, we can think of that node as contributing to many communities. As a clear example, consider the bottom two system labels in Fig. 4d, which represent nodes traditionally assigned to the visual system. Here, we find that edges associated with those visual nodes are spread out over the first four communities.

The overlapping nature of communities is made clearer in Fig. 4e, in which communities are represented topographically. The edges associated with the same visual nodes are all involved in communities 1, 2, 3, and 4 to some extent, thereby linking the visual system to multiple other brain systems. In community 1, for example, edges incident upon nodes in the visual and somatomotor systems are clustered together, whereas in community 3, edges are incident upon visual and default mode network nodes are assigned to the same community. There are other clear examples of this phenomenon involving other brain systems, including communities 8 and 9 and communities 5 and 10, which are dominated by edges incident upon the same somatomotor and default mode nodes, respectively.

Previous descriptions of the brain’s community structure have relied upon node-centric network representations and standard detection algorithms that force brain regions into non-overlapping communities [6, 11]. Even those approaches that have used overlapping methods to detect communities still relied on node-centric network representations, limiting the ability to draw inferences about the functional roles of network edges [46]. Viewed as a whole, these previous analyses reinforce a perspective in which brain function and community structure are both non-overlapping, with each community subtending a unique functional repertoire not shared by others. Our approach accommodates overlapping communities, corresponding to a view of brain function in which edge interactions engender polyfunctionality at the level of the node, naturally leading to functional roles that span traditional system boundaries.

### Sensorimotor systems participate in disproportionately many communities

In the previous section, we showed that the human cerebral cortex could be partitioned into overlapping communities based on its edge correlation structure. This observation leads to a series of additional questions. For instance, which brain areas participate in many communities? Which participate in few? If we changed the scale of inquiry – the number of detected communities – do the answers to these questions change? Do the answers depend on which dataset we analyze? In this section we explore these questions in detail.

At a coarse level, one strategy for assessing community overlap is to simply count the number of different communities to which each nodes’ edges are assigned [46]. These results, however, can be misleading – a brain region with 100 connections divided equally between two communities would have the same overlap as a brain region with 99 edges in one community and 1 edge in another. A more appropriate measure that accounts for the distribution of edge community assignments is a normalized entropy – a measurement that indexes the uniformity of a distribution. Intuitively, as the distribution of edge community assignments approaches uniformity its normalized entropy is close to 1; when edges are assigned to a single community normalized entropy is closer to 0 (See **Materials and Methods**). We calculated normalized entropy for every brain region while varying the number of communities from *k* = 2 to *k* = 20. In this section we focus on results with *k* = 10.

In general, we found that normalized entropy, a measure of community overlap, was greatest within sensorimotor and attentional systems, in contrast with previous reports [46]. In Fig. 5a,b,c we show results based on our analysis of HCP data (see Fig. S7 for corresponding plots using MSC and HBN datasets and Fig. S8 for normalized entropy maps at different values of *k*). Among the systems with the lowest normalized entropy were regions traditionally associated with control and default mode networks (Fig. 5c).

**FIG. 5.**
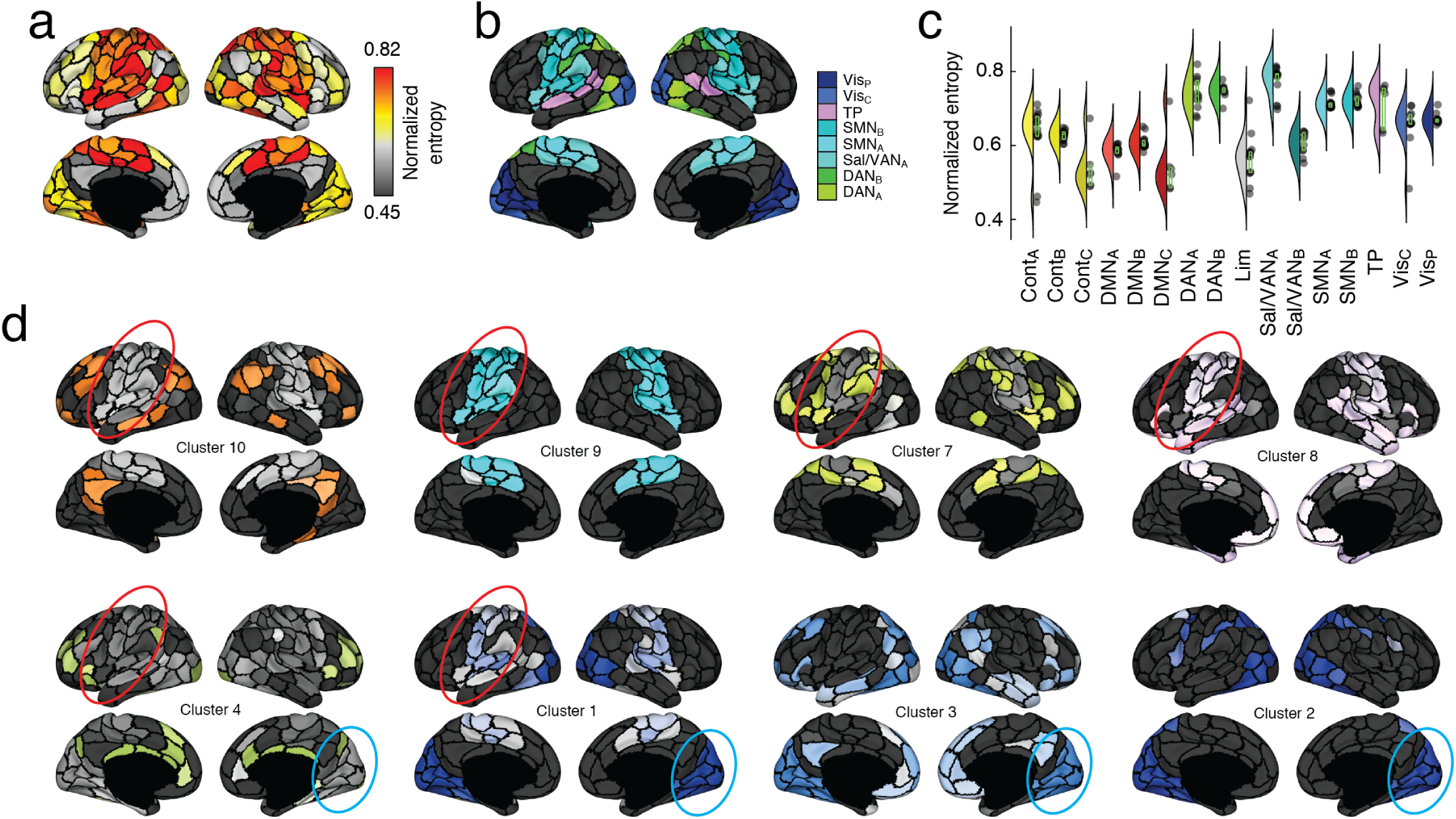
Edge community entropy and overlap. (*a*) Topographic distribution of normalized entropies. Normalized entropy, in this case, measures the uniformity of a node’s community assignments across all communities and serves as a measure of overlap. In general, higher entropy corresponds to greater levels of overlap. (*b*) Brain systems associated with the highest levels of normalized entropy. These include visual, attentional, somatmotor, and temporo-parietal systems. (*c*) Entropy values for all brain systems. (*d*) Here, we highlight communities in which somatomotor (red) and visual (blue) systems are represented.

Past studies that applied non-overlapping community detection algorithms to node-centric FC have used measures like the participation coefficient [47] to identify brain regions whose connectivity patterns span community boundaries [10, 39, 48, 49]. In particular, these studies have identified brain regions located within frontoparietal and cingulo-opercular association cortices as candidate “hubs.” Here, using an alternative method for defining overlapping communities, we find evidence of substantial overlap in sensorimotor and attentional system and less overlap in control and default mode systems. These observations suggest that, while control and default mode networks are relatively stable in their communication patterns, sensorimotor and attentional networks are flexible and can adapt their communication partners to meet ongoing cognitive demands. While the new methodology presented here can be immediately deployed and applied to investigate changes in overlapping community structure with cognitive, developmental, and disease state, understanding how overlapping communities reconfigure to form grander, higher-order structures in a network, e.g. cores and peripheries [37, 50, 51], remains an important open question for future studies.

### Edge functional connectivity is modulated by changes in sensory input

In the previous sections, we demonstrated that eFC is a reliable marker of an individual and that by clustering eFC we naturally obtain overlapping communities. We leveraged this final observation to demonstrate that primary sensory and attentional systems participate in disproportionately more communities than association cortices. While illuminating, these analyses were carried out using data collected during task-free conditions, i.e. at rest. Analogous to previous studies documenting the effect of task on nodal FC, we expect that eFC is also modulated by task load. In this section, we explore the effect of passive movie-watching on eFC.

To address this question, we analyzed fMRI data from the Healthy Brain Network Serial Scanning Initiative recorded during rest and while watching the movie “Raiders of the Lost Ark.” The movie data was divided into six successive scan sessions, which we further truncated by retaining the first 420 samples so that the duration matched that of the HBN resting scans, of which we retained the first six for the sake of balance. Then we estimated group-averaged eFC separately for each of the movie and rest scans.

In general, we found that eFC during movie-watching was highly correlated with eFC estimated during rest (Fig. 6a). Across all six movie scans, the mean correlation with resting eFC was *r* = 0.55±0.02. When we compared the pairwise similarity of all movie-watching scans with rest, we found that similarity of eFC was greater within a given task than between tasks (Fig. 6b). To better understand what was driving this effect, we computed the element-wise difference between the average movie and rest eFC matrices. While eFC differences were widespread, the most pronounced effects were associated with two specific communities (Fig. 6c), one of which exhibited a decrease in its within-module eFC, while they both decreased eFC with respect to each other. These communities, interestingly, consisted of edges associated with somatomotor and visual systems (Fig. 6d).

**FIG. 6.**
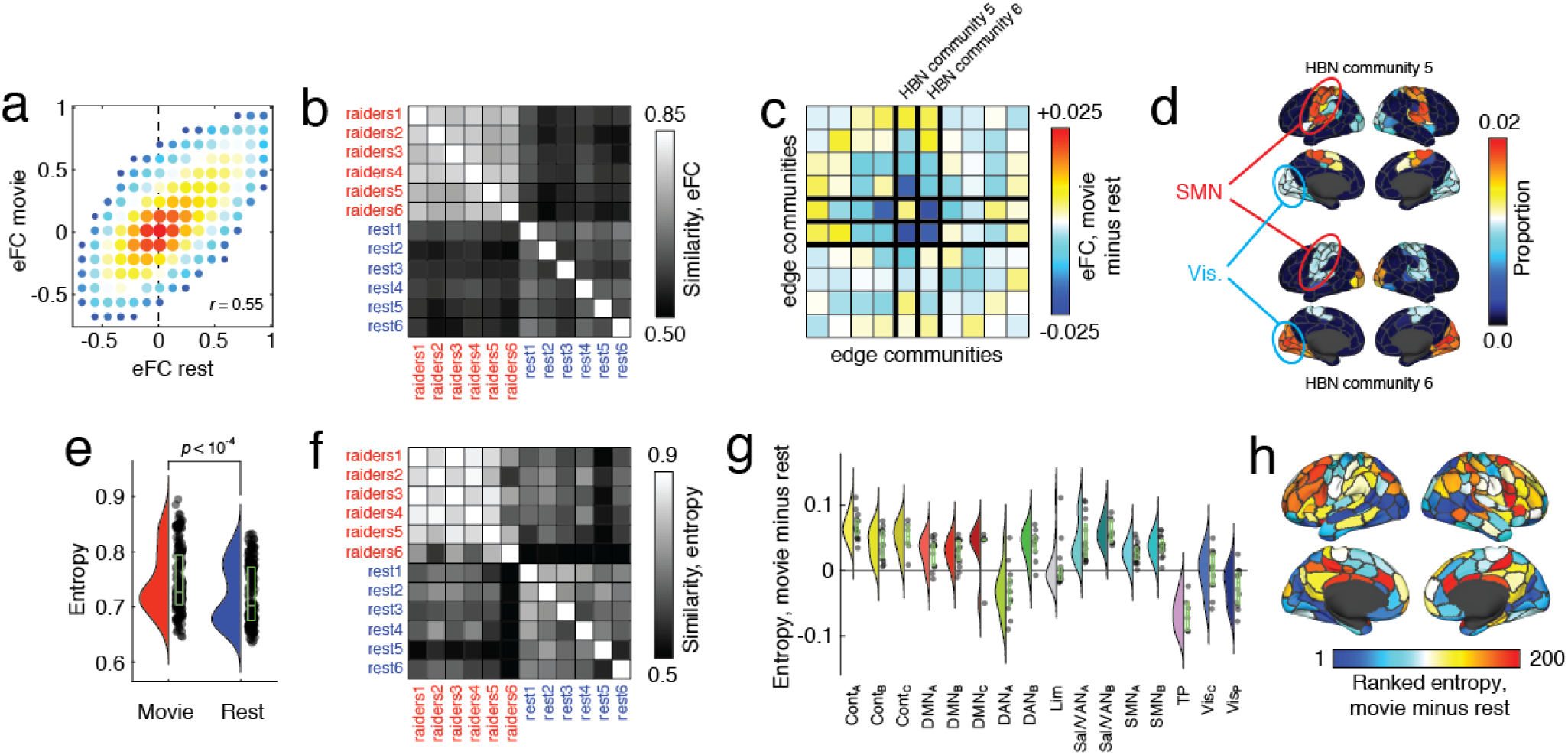
Effect of passive movie-watching on eFC. (*a*) Two-dimensional histogram of eFC estimated at rest with eFC estimated during movie-watching. (*b*) Similarity of whole-brain eFC estimated at rest with movie-watching. Note that within-condition similarity is greater for both conditions. (*c*) Community-averaged differences in eFC. Communities 5 and 6 are associated with the largest magnitude differences, on average. Note: these are communities estimated from HBN data and are not identical to those shown in Fig. 4 and estimated from HCP data. (*d*) Topographic distribution of communities 5 and 6. Note that these communities involve edges associated with visual and somatomotor systems. (*e*) Averaged differences in community overlap (normalized entropy). (*f*) Similarity of whole-brain normalized entropy estimated at rest with movie-watching. (*g*) Violin plot showing system-specific differences in normalized entropy. Note that some of the greatest increases in entropy are concentrated with control and default mode networks. (*h*) Topographic distribution of differences in entropy.

The differences in the connection weights of eFC between movie-watching and rest strongly suggested that the overall modular structure and the locations of high and low cluster overlap might also differ between conditions, as well. To test this, we used the same clustering algorithm described earlier to partition node pairs into non-overlapping clusters and, based on these clusters, calculated each node’s cluster overlap as a normalized entropy. We found that compared to rest, entropy increased during movie-watching (*p <* 0.01), indicating more overlap between communities (Fig. 6e), and that the brainwide pattern of entropy also differed (Fig. 6f). These differences could be attributed to increased entropy in association cortices, with control and default areas increasing by the greatest amount, on average (Fig. 6,g,h).

Collectively, these results suggest that, like nFC, eFC is reconfigurable and can be modulated by sensory inputs and ongoing task demands. The observed changes in eFC, which implicated two clusters associated both with somatomotor and visual systems, is in close agreement with past studies of passive movie-watching that documented changes in activity and nFC in similar areas [52, 53]. We also found increased overlap in areas associated with control and default mode networks, which agrees with other recent studies reporting that activity throughout these areas, and in default mode in particular, is sensitive to movie narrative structure [54] and comprehension [55]. An important area of future research involves systematically assessing the effect of different cognitively demanding tasks on eFC.

## DISCUSSION

Here, we presented a network model of human cerebral cortex that focused on edge-edge interactions. The network formed by these interactions, a construct we referred to as edge functional connectivity (eFC), was similar across datasets and more similar within subjects than between. When clustered, eFC provided a natural estimate of overlapping community structure. We found that the amount of overlap varied across the cortex, but peaked in sensorimotor and attention networks. Lastly, we showed that eFC and community overlap varied systematically during passive viewing of movies.

### Edge-centric perspective on functional network organization

Node-centric representations have dominated the field of network neuroscience and have served as the basis for nearly every discovery within that field [1]. As a result, alternative ways of representing functional neuroimaging data have not been thoroughly explored. The edge-centric representation shifts focus away from dyadic relationships between nodal activations and onto the interactions between edges (the similarity in their patterns of cofluctuation, a potential hallmark of communication), instead. While similar models have been explored in other scientific domains [19–22], they require as input sparse node-node connectivity matrices and are poorly suited for continuous-valued time series data, making them suboptimal representations of dynamic neural data.

Here, we developed a novel edge-centric representation of functional neuroimaging data that operates directly on observed time series. Our method for estimating connection weights between edges can be viewed as a temporal “unwrapping” of the familiar Pearson correlation – the measure frequently used to estimate the magnitude of nFC between pairs of brain regions [25]. Whereas the Pearson correlation coefficient calculates the time-averaged co-fluctuation magnitude for node pairs, we simply omit the averaging step, yielding “edge time series,” which represent the co-fluctuation magnitude between two nodes at every instant in time. This simple step enables us to track fluctuations in edge weight across time and, critically, can be related to one another dyadically, creating an entirely new network, edge-centric representation of nervous system architecture.

At its most basic level, edge time series are measuring instantaneous co-fluctuations between the activity of two brain regions. However, if we interpret edge time series as a temporal unwrapping of nFC, which is thought to reflect the aggregate effect of communication processes between neural elements [56, 57], then edge times series track, with unprecedented temporal resolution, the communication patterns between distributed neural elements.

### Overlapping communities extend our understanding of system-level cortical organization

Here, we demonstrated that clustering eFC using community detection methods naturally leads to communities that overlap when mapped back to the level of brain regions and nodes. Because eFC reflects interactions between edges, and because we interpret edge time series as time-resolved estimates of inter-regional communication, edge communities can be viewed as reflecting the brain’s circuit-level architecture. This is in contrast to nodal communities, which reflect similarity in regional activation patterns.

Past investigations of cortical organization have focused almost exclusively on non-overlapping communities [6, 11, 12, 38, 39]. The decision to define communities in this way is partially motivated by interpretability but also by limitations of the methods used to detect communities, which assign nodes to one community, only [44, 45] (although application of community detection methods to time-varying or multi-layer networks can overcome this to some extent [58–60]). This current view of communities has been profoundly successful. It provides a low-dimensional description of the brain [61], it can be used to define node roles and to detect hubs [47, 62], and can be applied to both anatomical [63] and functional networks [64] with equal success.

The dominant non-overlapping perspective of communities has strongly influenced we think about brain function. Because functional communities exhibit a correspondence with patterns of task-evoked activity [11, 65, 66], and because those patterns are stereotypical and reliably detected across multiple studies, we have come to associate individual communities with specific cognitive domains, labeling communities in reference to their supposed function. For instance, it is not uncommon to refer to communities as primarily processing visual information, enacting cognitive control, or performing attentional functions. This localization of brain function to communities, though likely a reasonable first-order approximation, perpetuates a view of brain function in which brain areas, systems, and communities are fundamentally unifunctional. Such a view, however, disagrees with observations that many aspects of cognition and behavior transcend these traditional community labels, such that the same neural substrate can be activated under different unrelated conditions.

The approach developed here is closely aligned with this second view, in which brain areas and communities exhibit highly degenerate functionality. Other studies have investigated overlapping communities in nFC, where communities represent clusters of activations [46, 67, 68]. Our approach, on the other hand, emphasizes clusters of communication patterns and offers a novel and complementary perspective on communities. While we apply edge-based clustering to functional MRI data, we note that this method is general and could be applied to virtually any neural time series data, irrespective of its provenance. Future work could pursue this possibility and investigate eFC in both scalp and intra-cranial EEG, MEG, electrophysiological recordings [3], as well as microscopy techniques [69], revealing novel organizational features of brain networks at different spatiotemporal scales.

### Limitations

In this study, we explore an edge-centric representation of human cerebral cortex. While this approach has provided new insight into cortical organization, there are several limitations worth discussing.

One of the most important limitations concerns the estimation of edge time series from functional imaging data. To calculate edge time series, we first z-score regional time series. Here, the z-score is only appropriate if the time series has a temporally invariant mean and standard deviation [70]. If there is a sustained increase or decrease in activity, e.g. the effect of a blocked task, then the z-scoring procedure can result in a biased mean and standard deviation, resulting in poor estimates of fluctuations in activity. To minimize the likelihood of this occurring, we focused on resting-state and movie-watching data rather than blocked tasks. In future work, investigation of task-evoked changes in eFC could be investigated with already common preprocessing steps, e.g. constructing task regressors to remove the first-order effect of tasks on activity [71].

Another limitation concerns the scalability of eFC. Calculating eFC given for a brain divided into *N* parcels results in an eFC matrix of dimensions *N*(*N* − 1)/2. This means that an increase in the number of parcels results in a squared increase in the dimensionality of eFC; if the number of parcels is large, then this can result in large, fully-weighted matrices that require large amounts of memory to store and manipulate. Here, we circumvented this issue by focusing on parcellations of the brain into 100-, 200-, and 400-node parcellations [33]. In the future, however, it may be necessary to explore dimension reduction methods to retain the most relevant dimensions for a given task or set of behaviors [72, 73].

### Future directions

While eFC characterizes interactions between edges rather than nodes, it can still be analyzed using the same methods previously applied to nFC. We can detect its hubs and communities [74], identify its central elements [75], estimate edge gradients [76], and compare eFC connection weights across individuals [36] and conditions [5]. On the other hand, eFC affords many new opportunities, beginning with the edge time series used to estimate eFC. Essentially, edge time series offer a moment-to-moment assessment of how strongly two nodes (brain regions) co-fluctuate with one another, providing an estimate of time-varying nFC without the requirement that we specify a window [77–79]. This overcomes one of the main limitations of sliding window estimates of time-varying nFC, namely that the use of a window leads to a “blurring” of events across time [80].

The eFC matrices, themselves, offer distinct advantages, some of which were explored here. For instance, partitions of eFC result in communities of edges, which can be easily mapped back to brain regions, thereby generating estimates of community overlap. We also note that, though not explored here, similar edge-to-node mappings can be performed using any network metric estimated on eFC, rendering edge statistics more interpretable and comparable to the more familiar network metrics for nFC [18]. While the present community analysis was carried out on parcel-level time series, a similar approach could be applied to downsampled surface-level time series to generate whole-brain functional atlases with overlapping system labels [6, 81] or applied to specific brain areas and sub-systems for constructing fine-grained overlapping atlases [82, 83].

eFC might also be particularly useful in applications of machine learning and classification of neuroimaging data [84, 85]. The dimensionality of the eFC matrix is much greater than that of a typical nFC matrix. Some of the added dimensions may be useful identifying manifolds along which subjects, clinical cohorts, or behaviors naturally separate, enhancing classification accuracy [86].

Lastly, the edge-centric framework developed here is not unique to functional MRI and can be easily be extended to different imaging modalities, including scalp/intracranial EEG or MEG, which making it possible to track seizure propagation at the level of edges [87]. Similar, the application of this approach to datasets resolving single neuron activity could uncover important connection-level insight into circuit organization [3].

## MATERIALS AND METHODS

### Datasets

The Human Connectome Project (HCP) dataset [26] included resting state functional data (rsfMRI) from 100 unrelated adult subjects (54% female, mean age = 29.11 ± 3.67, age range = 22-36). The study was approved by the Washington University Institutional Review Board and informed consent was obtained from all subjects. Subjects underwent four 15 minute rsfMRI scans over a two day span. A full description of the imaging parameters and image prepocessing can be found in [88]. The rsfMRI data was acquired with a gradient-echo EPI sequence (run duration = 14:33 min, TR = 720 ms, TE = 33.1 ms, flip angle = 52°, 2 mm isotropic voxel resolution, multiband factor = 8) with eyes open and instructions to fixate on a cross. Images were collected on a 3T Siemens Connectome Skyra with a 32-channel head coil.

The Midnight Scan Club (MSC) dataset [27] included rsfMRI from 10 adults (50% female, mean age = 29.1 ± 3.3, age range = 24-34). The study was approved by the Washington University School of Medicine Human Studies Committee and Institutional Review Board and informed consent was obtained from all subjects. Subjects underwent 12 scanning sessions on separate days, each session beginning at midnight. 10 rsfMRI scans per subject were collected with a gradient-echo EPI sequence (run duration = 30 min, TR = 2200 ms, TE = 27 ms, flip angle = 90°, 4 mm isotropic voxel resolution) with eyes open and with eye tracking recording to monitor for prolonged eye closure (to assess drowsiness). Images were collected on a 3T Siemens Trio.

The Healthy Brain Network Serial Scanning Initiative (HBN) dataset [28] included rsfMRI and movie watching (mwfMRI) data from 13 adults (54% female, mean age = 30.3 ± 6.4, age range = 21-42). Three subjects of the HBN dataset did not have enough non-outlier functional scans (see quality control criteria below) to be meaningfully analyzed (non outlier scan percentage = 7%, 0%, and 0%), and were excluded entirely from the current study. This rendered the HBN dataset as 10 subjects (50% female, mean age = 29.8 ± 5.3, age range = 23-37). The study was approved by the Chesapeake Institutional Review Board and informed consent was obtained from all subjects. Subjects underwent 14 scanning sessions over a 1-2 month period, in which 13 rsfMRI runs were acquired per subject. On the 8th session, subjects viewed the movie “Raiders of the Lost Ark” (Lucasfilm Ltd., 1981) in six approximately 20 minute scans. The rsfMRI and mvfMRI were acquired with a gradient-echo EPI sequence (run duration rsfMRI = 10:18 min, mvfMRI = 20 min per segment, TR = 1450 ms, TE = 40 ms, flip angle = 55°, 2.46×2.46×2.5 mm voxel resolution, multiband factor = 3) with subjects instructed to keep their eyes open and gazed directed towards a cross during the fsMRI scan. Images were collected on a 1.5T Siemens Avanto with a 32-channel head coil.

### Image Preprocessing

#### HCP Functional Preprocessing

Functional images in the HCP dataset were minimally preprocessed according to the description provided in [88]. Briefly, these data were corrected for gradient distortion, susceptibility distortion, and motion, and then aligned to a corresponding T1-weighted (T1w) image with one spline interpolation step. This volume was further corrected for intensity bias and normalized to a mean of 10000. This volume was then projected to the *32k_fs_LR* mesh, excluding outliers, and aligned to a common space using a multi-modal surface registration [89]. The resultant cifti file for each HCP subject used in this study followed the file naming pattern: `*_REST{1,2} {L,R} Atlas MSMAll.dtseries.nii.`

#### MSC and HBN Functional Preprocessing

Functional images in the MSC and HBN datasets were preprocessed using *fMRIPrep* 1.3.2 [90], which is based on Nipype 1.1.9 [91]. The following description of *fMRIPrep*’s preprocessing is based on boilerplate distributed with the software covered by a “no rights reserved” (`CC0`) license. Internal operations of *fMRIPrep* use Nilearn 0.5.0 [92], ANTs 2.2.0, FreeSurfer 6.0.1, FSL 5.0.9, and AFNI v16.2.07. For more details about the pipeline, see the section corresponding to workflows in *fMRIPrep*’s documentation.

The T1-weighted (T1w) image was corrected for intensity non-uniformity with `N4BiasFieldCorrection` [93, 94], distributed with ANTs, and used as T1w-reference throughout the workflow. The T1w-reference was then skull-stripped with a Nipype implementation of the `antsBrainExtraction.sh` workflow, using NKI as the target template. Brain surfaces were reconstructed using `recon-all` [95], and the brain mask estimated previously was refined with a custom variation of the method to reconcile ANTs-derived and FreeSurfer-derived segmentations of the cortical gray-matter using Mindboggle [96]. Spatial normalization to the *ICBM 152 Nonlinear Asymmetrical template version 2009c* [97] was performed through nonlinear registration with `antsRegistration`, using brain-extracted versions of both T1w volume and template. Brain tissue segmentation of cerebrospinal fluid (CSF), white-matter (WM) and gray-matter (GM) was performed on the brain-extracted T1w using FSL’s `fast` [98].

Functional data was slice time corrected using AFNI’s `3dTshift` and motion corrected using FSL’s `mcflirt` [99]. *Fieldmap-less* distortion correction was performed by co-registering the functional image to the same-subject T1w image with intensity inverted [100] constrained with an average fieldmap template [101], implemented with `antsRegistration`. This was followed by co-registration to the corresponding T1w using boundary-based registration [102] with 9 degrees of freedom. Motion correcting transformations, field distortion correcting warp, BOLD-to-T1w transformation and T1w-to-template (MNI) warp were concatenated and applied in a single step using `antsApplyTransforms` using Lanczos interpolation. Several confounding time-series were calculated based on this preprocessed BOLD: framewise displacement (FD), DVARS and three region-wise global signals. FD and DVARS are calculated for each functional run, both using their implementations in Nipype [103]. The three global signals are extracted within the CSF, the WM, and the whole-brain masks. The resultant nifti file for each MSC and HBN subject used in this study followed the file naming pattern `*_space-T1w desc-preproc bold.nii.gz.`

### Image Quality Control

All functional images in the HCP dataset were retained. The quality of functional images in the MSC and HBN were assessed using *fMRIPrep*’s visual reports and *MRIQC* 0.15.1 [104]. Data was visually inspected for whole brain field of view coverage, signal artifacts, and proper alignment to the corresponding anatomical image. Functional data were excluded if greater than 25% of the frames exceeded 0.2 mm framewise displacement [105]. Furthermore, functional data were excluded if marked as an outlier (exceeding 1.5x inter-quartile range in the adverse direction) in more than half of the following image quality metrics (calculated within-dataset, across all functional acquisitions): *dvars, tsnr, fd mean, aor, aqi, snr*, and *efc*. Information about these image quality metrics can be found within *MRIQC* ‘s documentation [106].

### Functional and Structural Networks Preprocessing

#### Parcellation Preprocessing

A functional parcellation designed to optimize both local gradient and global similarity measures of the fMRI signal [33] (*Schaefer200*) was used to define 200 areas on the cerebral cortex. These nodes are also mapped to the Yeo canonical functional networks [6]. For the HCP dataset, the *Schaefer200* is openly available in *32k_fs_LR* space as a cifti file. For the MSC and HBN datasets, a *Schaefer200* parcellation was obtained for each subject using a Gaussian classifier surface atlas [107] (trained on 100 unrelated HCP subjects) and FreeSurfer’s `mris_ca_label` function. These tools utilize the surface registrations computed in the `recon-all` pipeline to transfer a group average atlas to subject space based on individual surface curvature and sulcal patterns. This method rendered a T1w space volume for each subject. For use with functional data, the parcellation was resampled to 2mm T1w space. This process could repeated for other resolutions of the parcellation (i.e. *Schaefer100*).

#### Functional Network Preprocessing

The mean BOLD signal for each cortical node data was linearly detrended, band-pass filtered (0.008-0.08 Hz) [105], confound regressed and standardized using Nilearn’s `signal.clean`, which removes confounds orthogonally to the temporal filters [108]. The confound regression employed [109] included 6 motion estimates, time series of the mean CSF, mean WM, and mean global signal, the derivatives of these nine regressors, and the squares these 18 terms. Furthermore, a spike regressor was added for each fMRI frame exceeding a motion threshold (HCP = 0.25 mm root mean squared displacement, MSC, HBN = 0.5 mm framewise displacement). This confound strategy has been shown to be relatively effective option for reducing motion-related artifacts [105]. Following this preprocessing and nuisance regression, residual mean BOLD time series at each node was recovered. eFC matrices for each subject were computed and then averaged across subjects, to obtain a representative eFC matrix for each dataset. This processing was performed for both resting state and movie watching data.

### Edge graph construction

Constructing networks from fMRI data (or any neural time series data) requires estimating the statistical dependency between every pair of time series. The magnitude of that dependency is usually interpreted as a measure of how strongly (or weakly) those voxels are parcels are functionally connected to each other. By far the most common measure of statistic dependence is the Pearson correlation coefficient. Let **x**_*i*_ = [*x*_*i*_(1), …, *x*_*i*_(*T*)] and **x**_*j*_ = [*x*_*j*_(1), …, *x*_*j*_(*T*)] be the time series recorded from voxels or parcels *i* and *j*, respectively. We can calculate the correlation of *i* and *j* by first z-scoring each time series, such that 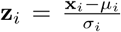, where 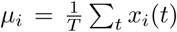 and 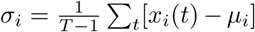 are the time-averaged mean and standard deviation. Then, the correlation of *i* with *j* can be calculated as: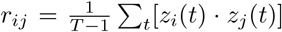. Repeating this procedure for all pairs of parcels results in a node-by-node correlation matrix, i.e. an estimate of FC. If there are *N* nodes, this matrix has dimensions [*N* × *N*].

To estimate *edge*-centric networks, we need to modify the above approach in one small but crucial way. Suppose we have two z-scored parcel time series, **z**_*i*_ and **z**_*j*_. To estimate their correlation we calculate the mean their element-wise product (not exactly the average, because we divide by *T* − 1 rather than *T*). Suppose, instead, that we never calculate the mean and simply stop after calculating the element-wise product. This operation would result in a vector of length *T* whose elements encode the moment-by-moment co-fluctuations magnitude of parcels *i* and *j*. For instance, suppose at time *t*, parcels *i* and *j* simultaneously increased their activity relative to baseline. These increases are encoded in **z**_*i*_ and **z**_*j*_ as positive entries in the *t*th position, so their product is also positive. The same would be true if *i* and *j decreased* their activity simultaneously (because the product of negatives is a positive). On the other hand, if *i* increased while *j* decreased (or *vice versa*), this would manifest as a negative entry. Similarly, if either *i* or *j* increased or decreased while the activity of the other was close to baseline, the corresponding entry would be close to zero.

Accordingly, the vector resulting from the elementwise product of **z**_*i*_ and **z**_*j*_ can be viewed as encoding the magnitude of moment-to-moment co-fluctuations between *i* and *j*. An analogous vector can easily be calculated for every pair of parcels (network nodes), resulting in a set of co-fluctuation (edge) time series. With *N* parcels, this results in 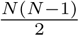 pairs, each of length *T*. From these time series we can estimate the statistical dependency for every pair of edges. We refer to this construct as edge functional connectivity (eFC). Let **c**_*ij*_ = [*z*_*i*_(1) · *z*_*j*_(1), …, *z*_*i*_(*T*) · *z*_*j*_(*T*)] and **c**_*uv*_ = [*z*_*u*_(1)·*z*_*v*_(1), …, *z*_*i*_(*T*)·*z*_*j*_(*T*)] be the time series for edges {*i, j*} and {*u, v*}, respectively. Then we can calculate eFC as:

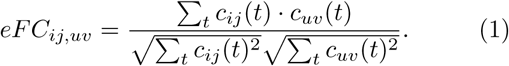

Here, the denominator is necessary to bound eFC to the interval [−1, 1].

### Clustering algorithm

In general, eFC matrices are much larger than traditional nodal FC matrices. While most clustering algorithms can be applied to hundreds or even thousands of observations, estimating clusters for eFC (which consists of tens of thousands of observations, each paired with at least as many features), presents a computational challenge, especially if the aim is to explore the space of possible partitions. To address this issue and to cluster eFC, we developed a simple two-step clustering procedure that operates on a low-dimensional representation of the eFC matrix.

First, we performed an eigendecomposition of the eFC matrix, retaining the top 50 eigenvectors. These eigenvectors were rescaled to the interval [-1, 1] by dividing each eigenvector by its largest magnitude element. Then simply clustered the rescaled eigenvectors using a standard k-means algorithm with Euclidean distance. We varied the number of communities, *k*, from *k* = 2 to *k* = 20, repeating the clustering algorithm 250 at each value. We retained as a representative partition the one with the greatest overall similarity to all other partitions. We note that the edge time series can be clustered directly and that, in general, the results were highly similar (Fig. S9).

### Community overlap metrics

The clustering algorithm partitioned edges into non-overlapping clusters. That is, every edge {*i, j*}, where *i, j* ∈ {1, …, *N*}, was assigned to one of *k* clusters. In this list of edges, each node appeared *N* − 1 times (we excluded self-connections). Region *i*’s participation in cluster *c* was calculated as:

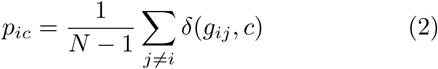

where *g*_*ij*_ ∈ {1, …, *k*} was the cluster assignment of the edge linking nodes *i* and *j* and *δ*(*x, y*) is the Kronecker delta, whose value is 1 if *x* = *y* and zero otherwise.

By definition, Σ_*c*_*pic* = 1, and we can treat the vector **p**_*i*_ = [*p* _*i*1_, …, *p*_*ik*_] as a probability distribution. The entropy of this distribution measures the extent to which region *i*’s community affiliations are distributed evenly across all communities (high entropy and high overlap) or concentrated within a small number of communities (low entropy and low overlap). We calculate this entropy as:

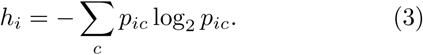

To normalize this measure and bound it to the interval [0, 1], we divide by log_2_*k*. We refer to this measure as community entropy and interpret this value as an index of overlap.

## AUTHOR CONTRIBUTIONS

JF and RFB conceived of study, processed data, carried out all analyses, and wrote the first draft of the manuscript. FZE, YJ, and OS contributed to project direction *via* discussion. All authors helped revise and write the submitted manuscript.

## DATA AVAILABILITY

All imaging data come from publicly-available, open-access repositories. Human connectome project data can be accessed *via* https://db.humanconnectome.org/app/template/Login.vm after signing a data use agreement. Midnight scan club data can be accessed *via* OpenfMRI at https://www.openfmri.org/dataset/ds000224/ and the Healthy Brain Network Serial Scanning Initiative data can be accessed *via* http://fcon_1000.projects.nitrc.org/indi/hbn_ssi/download.html.

## CODE AVAILABILITY

All processing and analysis code is available upon reasonable request.

---

**FIG. S1.**
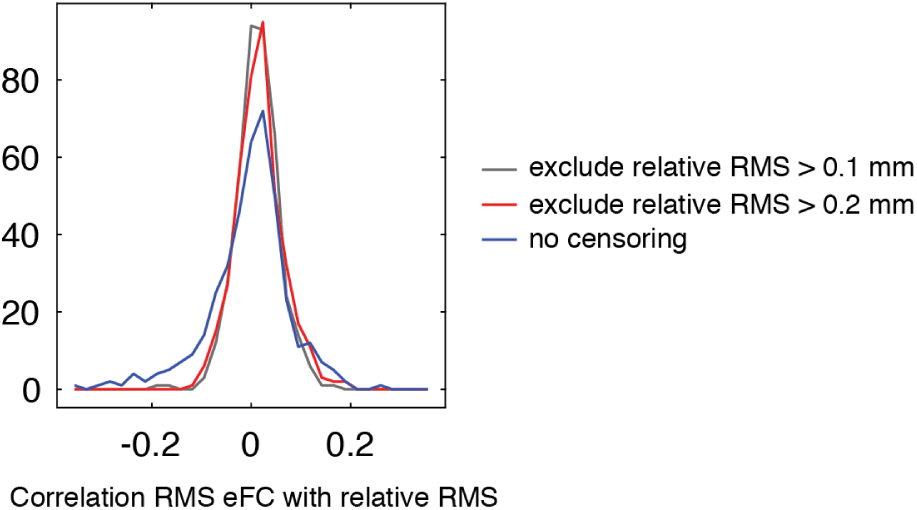
Effect of in-scanner motion on co-fluctuation time series. The edge time series shown Fig. 4a is typical and consistent with what we observed in other subjects. One of the most salient features of the edge time series is their sparse temporal structure. Rather the exhibiting smooth and continuous fluctuations over time, the edge time series exhibit long periods of quietude punctuated by large “events” when many edges simultaneously exhibit large shifts in amplitude. This pattern is not unlike typical framewise displacement plots, where excessive head motion is rarely sustained over long periods of time. This raises the concern that edge time series events are simple reflections of in-scanner head motion. To address this concern, we calculated the correlation of event amplitude at each time point (RMS eFC) with the instantaneous estimates of relative motion RMS. If the events were associated with in-scanner motion, we would expect the correlation to be strong and positive. We performed this procedure for all 100 subjects in the HCP dataset and for each of the four scans, independently. We carried out three versions of this analysis. In the first (labeled “no censoring”), we retained all volumes. In the other two we excluded volumes for which motion was greater than 0.1 mm and 0.2 mm, respectively. We found that, in general, the correlation of edge amplitude motion was weak. Without censoring, the median correlation across all 400 HCP scans was 0.008 with an interquartile range of [−0.039, 0.038]. With censoring, this distribution was narrowed slightly, so that the median value was still close to zero (0.01 in the case of censoring based on 0.1 mm and 0.2 mm) but with interquartile ranges of [−0.014, 0.045] and [−0.012, 0.041]. Collectively, these results suggest that the observed events not direct consequences of in-scanner motion.

**FIG. S2.**
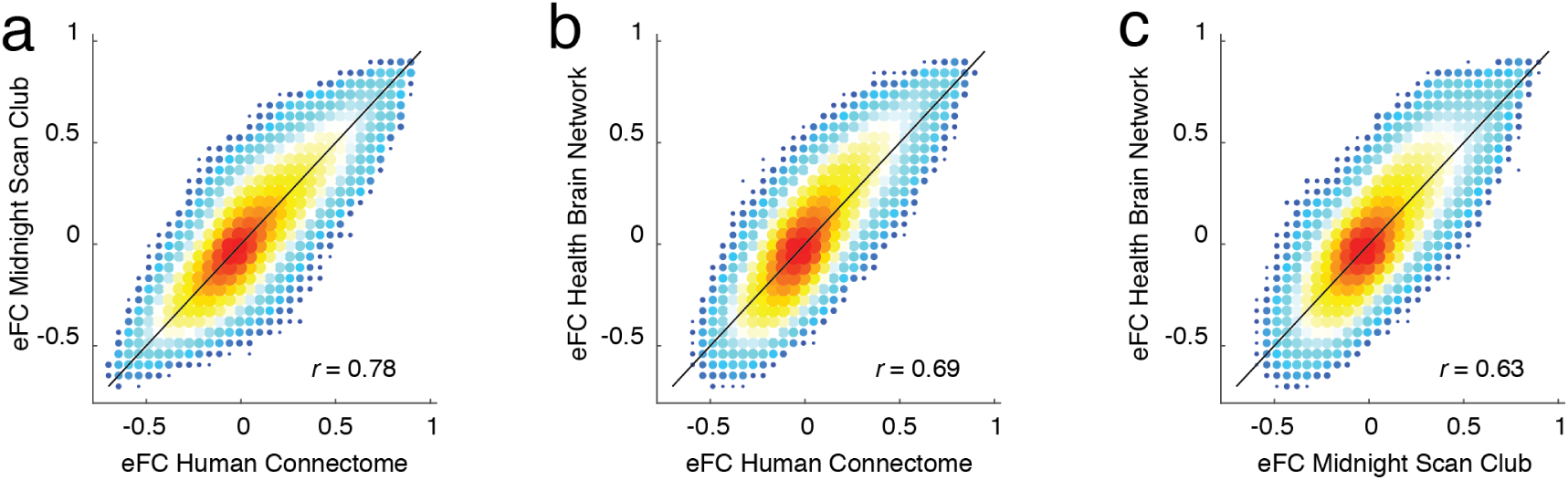
Comparison of group-averaged eFC across datasets. Here, we show two-dimensional histograms comparing the similarity of group-averaged eFC matrices for each of the three datasets: Human Connectome Project, Midnight Scan Club, and the Healthy Brain Network Serial Scanning Initiative.

**FIG. S3.**
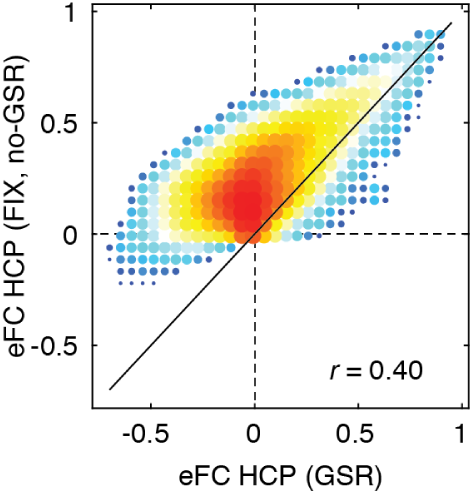
Effect of global signal regression on eFC. Here, we show two-dimensional histograms comparing the similarity of group-averaged eFC from the HCP dataset preprocessed with global signal regression (36 parameter) and without (ICA-FIX).

**FIG. S4.**
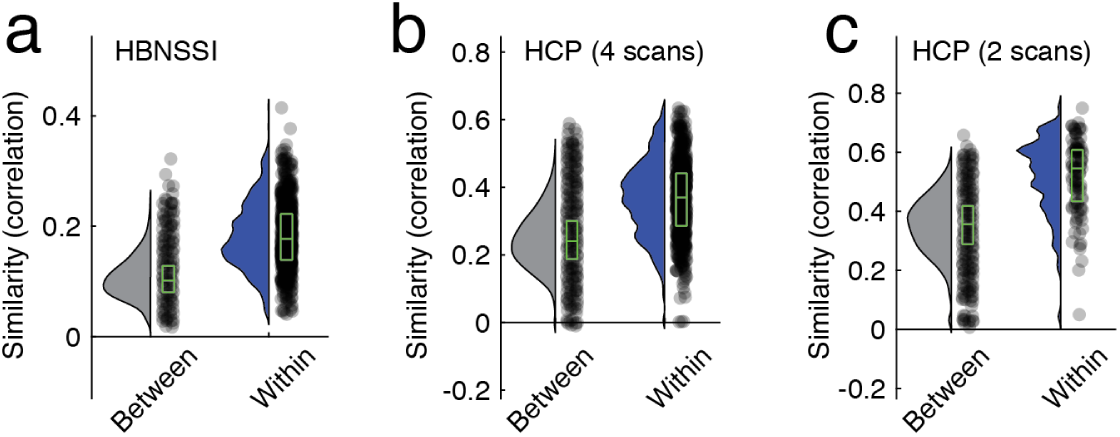
Within-versus between-subject similarity of eFC in HBN and HCP datasets. In the main text we showed, using data from the Midnight Scan Club, that eFC was more similar within subjects than between subjects. Here, we show analogous plots for the other two datasets: the Health Brain Network Serial Scanning Initiative and the Human Connectome Project. Panel *a* shows between- and within-subject similarity for the HBN dataset. Panels *b* and *c* depict an analogous figure for HCP data. Panel *b* calculates between- and within-subject similarity treating all four scans (REST1_LR, REST1_RL, REST2_LR, and REST2_RL) independently, whereas panel *c* shows similarity measures for combined REST1_LR + REST1_RL and REST2_LR + REST2_RL. Note that in all cases, the differences of between- and within-subjects similarity were statistically significant (maximum *p*-value of *p* ≈ 10^−6^)

**FIG. S5.**
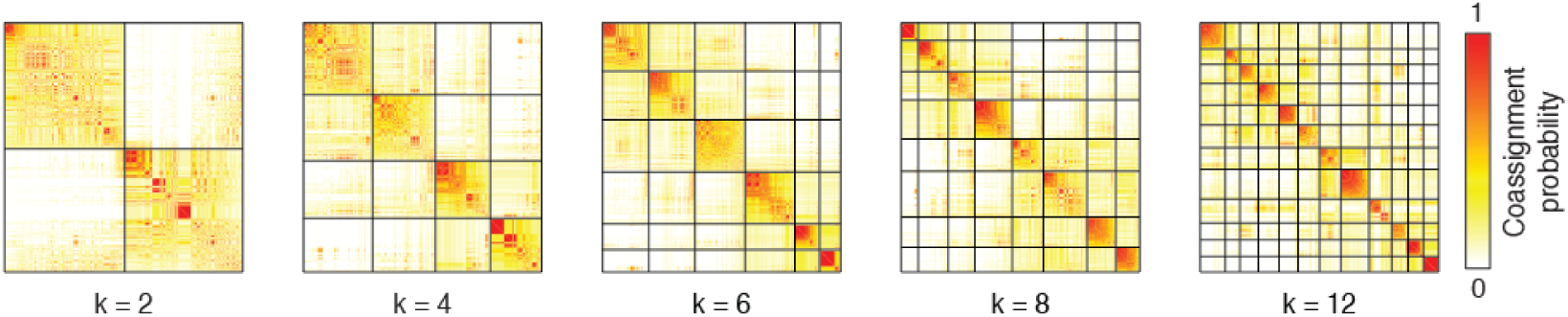
Co-assignment probability at different values of *k*. In the main text we showed the the matrix of co-assignment probabilities. In that figure, we reordered the rows and columns to highlight communities detected with *k* = 10. To do so, we truncated the upper limit on the co-assignment probability matrix to correspond with the *k* = 10 communities. Here, we show the same matrix with *k* = 2, *k* = 4, *k* = 6, *k* = 8, and *k* = 12 and without a truncated upper limit.

**FIG. S6.**
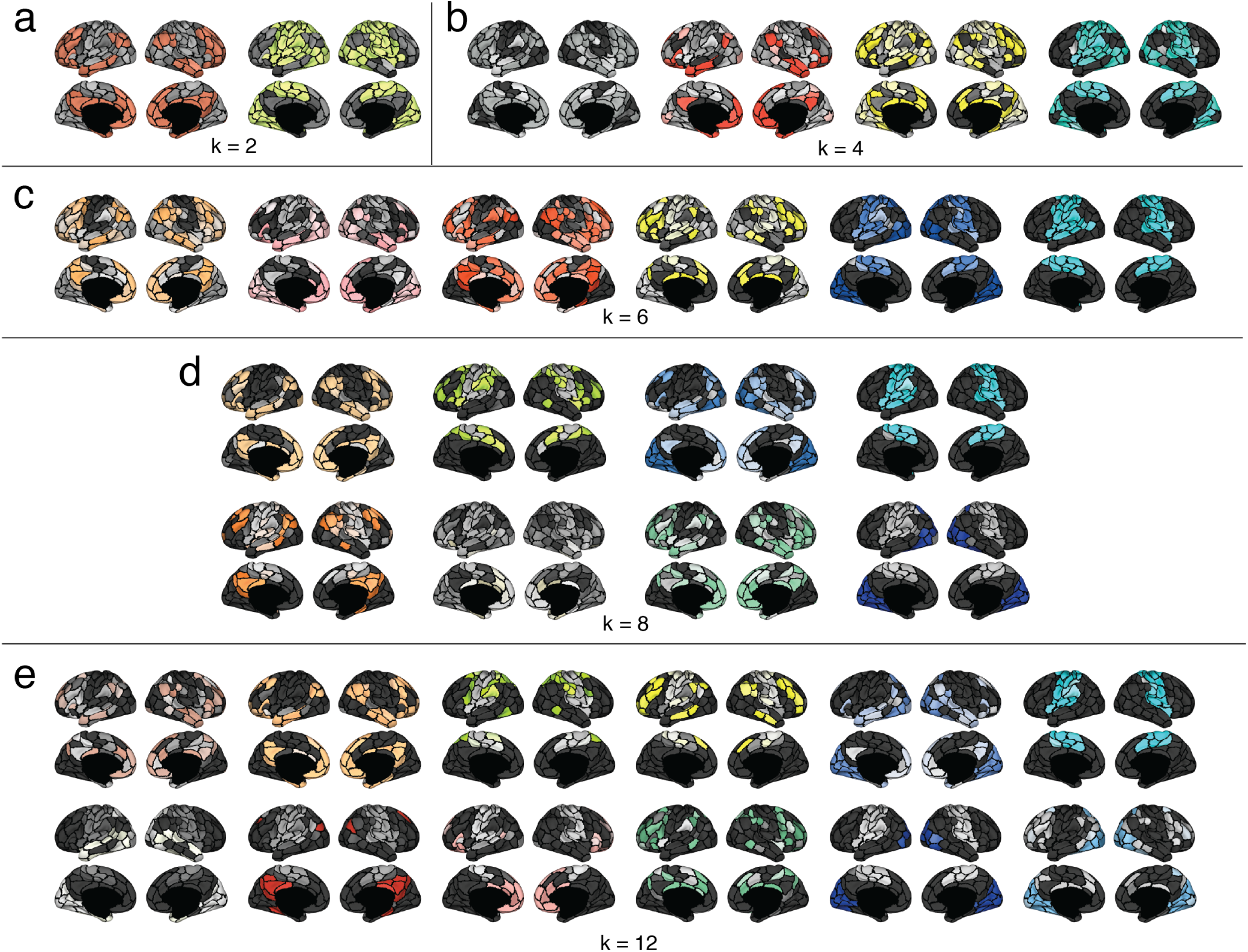
Effect of varying *k* on cluster structure. In the main text we detected clusters when the number of clusters was fixed at *k* = 10. Here, we show examples of clusters detected when *k* = 2, *k* = 4, *k* = 6, *k* = 8, and *k* = 12.

**FIG. S7.**
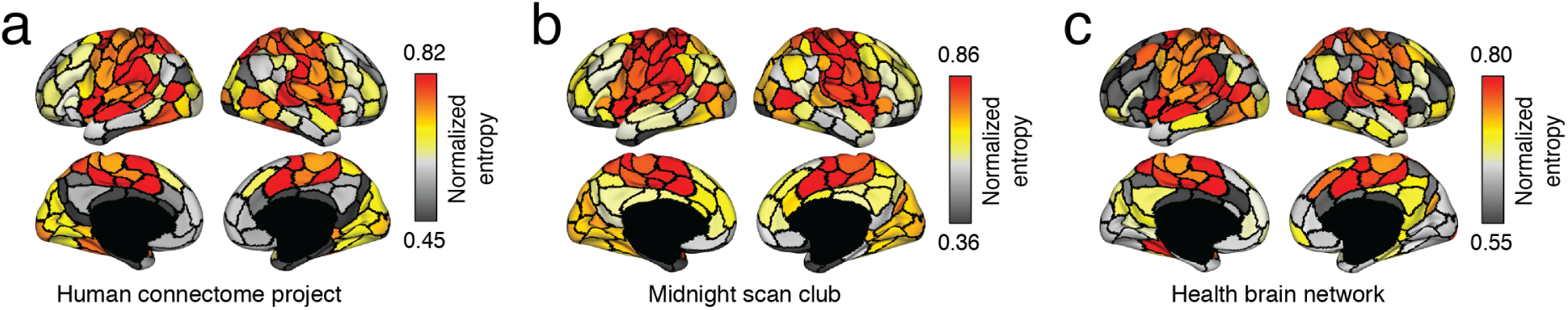
Comparison of normalized entropy across datasets. In the main text, we showed the normalized entropy – a measure of cluster overlap – for the HCP dataset. Here, we show the same measure for all three datasets. All plots correspond to *k* = 10 clusters.

**FIG. S8.**
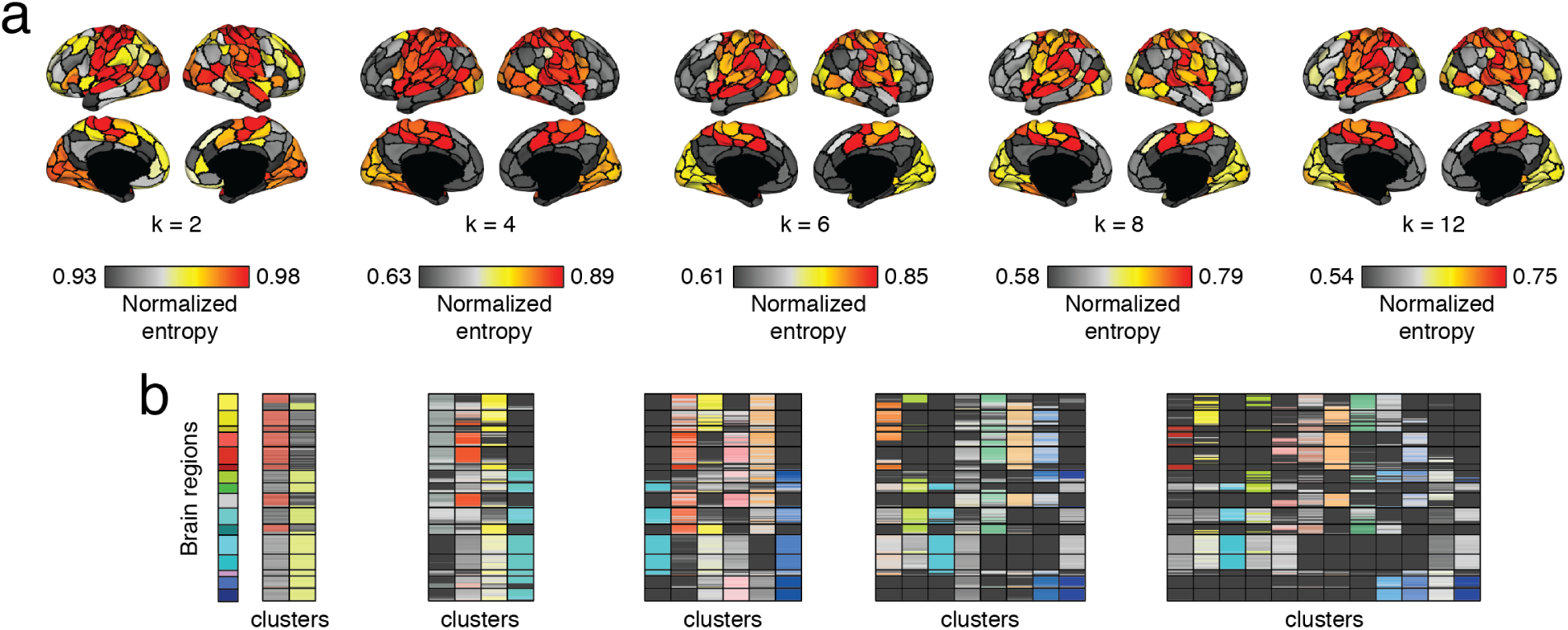
Effect of varying *k* on cluster overlap. In the main text, we showed normalized entropy with the number of clusters fixed at *k* = 10. In panel *a*, we show the same metric as we vary the number of clusters from *k* = 2, *k* = 4, *k* = 6, *k* = 8, and *k* = 12. Panel *b* shows matrix representations of the cluster assignments used to estimate normalized entropy.

**FIG. S9.**
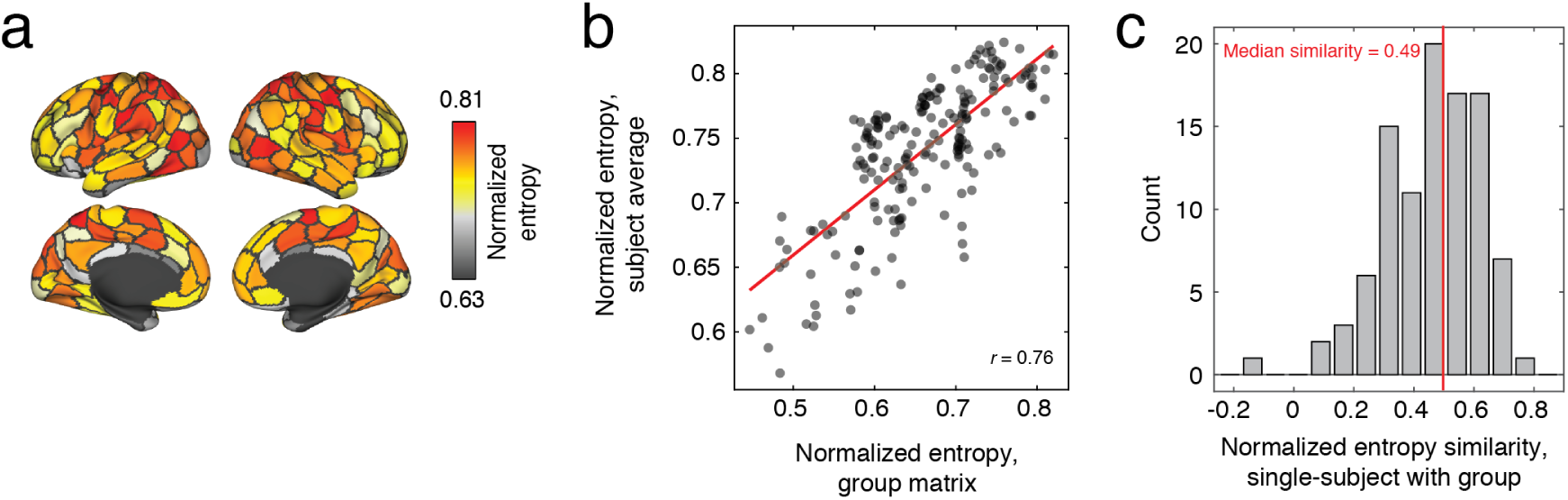
Comparison of normalized entropy from group-averaged data with single-subject data. In the main text, we showed the normalized entropy – a measure of cluster overlap – for the HCP dataset estimated by clustering a group-averaged eFC matrix that had undergone a dimension reduction step. Here, we show the similarity of that entropy pattern with entropies obtained from complete (i.e. no dimension reduction) single-subject data from the HCP dataset. (*a*) Mean normalized entropy pattern after averaging subject-level entropies. (*b*) Scatterplot showing the relationship of the subject average normalized entropy with the group-representative normalized entropy estimated in the main text. (*c*) Correlation coefficients (similarity) of single-subject entropies with the group-representative pattern described in the main text. The median similarity across all subjects was a correlation of *r* = 0.49.

